# Multiplex PCR with Nanopore Sequencing for Sequence-Based Detection of Four Tilapia Pathogens

**DOI:** 10.1101/2023.05.13.540096

**Authors:** Jérôme Delamare-Deboutteville, Watcharachai Meemetta, Khaettareeya Pimsannil, Han Ming Gan, Laura Khor Li Imm, Chadag Vishnumurthy Mohan, Ha Thanh Dong, Saengchan Senapin

**Affiliations:** WorldFish, Penang, Malaysia; Fish Health Platform, Center of Excellence for Shrimp Molecular Biology and Biotechnology (Centex Shrimp), Faculty of Science, Mahidol University, Bangkok, Thailand; Patriot Biotech Sdn Bhd, Bandar Sunway, 47500, Selangor, Malaysia; School of Environment, Resources and Development, Asian Institute of Technology, Pathum Thani, 12120, Thailand; National Center for Genetic Engineering and Biotechnology (BIOTEC), National Science and Technology Development Agency (NSTDA), Pathum Thani, Thailand

## Abstract

**Background:** Tilapia aquaculture faces significant threats posed by four prominent pathogens: tilapia lake virus (TiLV), infectious spleen and kidney necrosis virus (ISKNV), *Francisella orientalis*, and *Streptococcus agalactiae*. Currently, employed molecular diagnostic methods for these pathogens rely on multiple singleplex PCR reactions, which are time-consuming and expensive.

**Methods:** In this study, we present an approach utilizing a multiplex PCR (mPCR) assay, coupled with rapid Nanopore sequencing, enabling the one-tube simultaneous detection and one-reaction Nanopore sequencing-based validation of four pathogens.

**Results:** Our one-tube multiplex assay exhibits a detection limit of 1,000 copies per reaction for TiLV, ISKNV, and *S. agalactiae*, while for *F. orientalis*, the detection limit is 10,000 copies per reaction. This sensitivity is sufficient for diagnosing infections and co-infections in clinical samples from sick fish, enabling rapid confirmation of the presence of pathogens. Integrating multiplex PCR and Nanopore sequencing provides an alternative approach platform for fast and precise diagnostics of major tilapia pathogens in clinically sick animals, adding to the available toolbox for disease diagnostics.

## Introduction

Tilapia (*Oreochromis* spp.) is one of the most widely farmed freshwater fish species globally due to its high adaptability, fast growth, and excellent meat quality. Global production is estimated at 6,100,719 tonnes in 2020 (FAO, 2022). By providing sustenance, employment opportunities, and domestic and export revenues, this valuable species supports large populations worldwide (Wang & Lu, 2016). In the past decade, tilapia production has almost doubled (FAO, 2020), attributed mainly to its ease of cultivation, market demand, and stable pricing (Wang & Lu, 2016).

However, the production of tilapia has been threatened by several viral and bacterial diseases that can cause significant economic losses (Debnath et al., 2023). Among the most important tilapia pathogens are tilapia lake virus (TiLV), infectious spleen and kidney necrosis virus (ISKNV), *Francisella orientalis*, and *Streptococcus agalactiae* (SAG) also called group B *streptococcus* (GBS) (Kawasaki et al., 2018; Machimbirike et al., 2019; Mabrok et al., 2021; Haenen et al., 2023; Alathari et al., 2023). Effective diagnosis and monitoring of these pathogens are crucial for disease management and control (Dong et al., 2023). Traditional diagnostic techniques such as virus and bacterial isolation, immunofluorescence assay (IFA), and enzyme-linked immunosorbent assay (ELISA) are time-consuming, labor-intensive, and require specialized equipment and expertise (Dong et al., 2023). In contrast, molecular methods such as polymerase chain reaction (PCR) and quantitative PCR (qPCR) have gained widespread use in pathogen detection due to their high sensitivity and specificity, rapid turnaround time, and ability to detect multiple pathogens in a single reaction (Soto et al., 2010; Kralik & Ricchi, 2017; Liamnimitr et al., 2018; Waiyamitra et al., 2018; López□Porras et al., 2019; Ramírez-Paredes et al., 2021; Taengphu et al., 2022; Dong et al., 2023).

In current real-world situations, it is uncommon to perform subsequent sequencing of the positive PCR or qPCR products to validate the pathogen identity due to the time and resources involved in conventional Sanger sequencing. Sequence-based verification is particularly valuable in multiplex PCR since non-specific amplification is more likely to occur in a multiplex PCR reaction due to multiple primer combinations. Fortunately, this is now possible with Oxford nanopore technology (ONT) that allows on-site amplicon sequencing (Delamare-Deboutteville et al., 2021). Therefore, proper integration of multiplex PCR and ONT-based genotyping may present a promising tool for the efficient and precise detection and characterization of pathogens in tilapia aquaculture.

We present a multiplex PCR assay with a detection level suitable for simultaneously identifying TiLV, ISKNV, *F. orientalis*, and *S. agalactiae* in sick tilapia samples. Using the portable nanopore sequencing platform, we confirmed the presence of these pathogens at the sequence level and gathered genetic information from the amplicons. This assay offers a rapid and practical solution for detecting and genetically characterizing these pathogens, with the potential to complement existing diagnostic tools and inform targeted surveillance and control strategies.

## Materials & Methods

### Ethics declarations

The authors confirm that the journal’s ethical policies, as noted on the journal’s author guidelines page, have been adhered to. No ethical approval was required as no animals were used in this study. Virus sequences were generated from archived samples.

### Clinical samples and nucleic acid extraction

We utilized archival clinical samples of fry, fingerling, juvenile and adult Nile tilapia, red tilapia, and Asian sea bass from challenge experiments or from various disease outbreaks between 2015 and 2020. Our investigation included samples that were either confirmed to be caused by a single pathogen using PCR diagnosis or suspected to have resulted from co-infections. All sample details are comprehensively listed in Table 1. To extract the nucleic acid from the samples, some were processed using the PathoGen-spin DNA/RNA extraction kit from iNtRON Biotechnology, while others were archival RNA samples extracted using Trizol reagent from Invitrogen, and DNA samples extracted using the conventional phenol/chloroform ethanol precipitation method as described by (Meemetta et al., 2020). If done close to the farm, the diagnostic workflow from the point of sample collection to the final data analysis can take less than 12 hours. The processes include nucleic acid extraction, multiplex PCR, library preparation, Nanopore sequencing, and data analysis, as illustrated in Figure 1.

**Table 1.** Sources of clinical tilapia samples and PCR detection results.

| Code | Sample_name | Life stage | Country | Year Isolation | Previous Diagnosis sPCR | mPCR | Refs/comments |
| --- | --- | --- | --- | --- | --- | --- | --- |
| 1 | Healthy NT 1 | juvenile (NT) | Thailand | 2020 | - | - | 16 |
| 2 | Healthy NT 2 | juvenile (NT) | Thailand | 2020 | - | - | 16 |
| 3 | *r2_NT F1* | fingerling (NT) | Thailand | 2020 | TiLV (++) | TiLV* | 16 |
| 4 | *r2_NT F2* | fingerling (NT) | Thailand | 2020 | TiLV (++) | TiLV* | 16 |
| 5 | *r2_NT F3* | fingerling (NT) | Thailand | 2020 | TiLV (++) | TiLV* | 16 |
| 6 | *r2_NT F4* | fingerling (NT) | Thailand | 2020 | TiLV (++) | TiLV* | 16 |
| 7 | *r2_NT F5* | fingerling (NT) | Thailand | 2020 | TiLV (++) | TiLV* | 16 |
| 8 | NT fingerling pond C1 | fingerling (NT) | Thailand | 2020 | TiLV (+) | - | 16 |
| 9 | NT fingerling pond C2 | fingerling (NT) | Thailand |  | TiLV (+) | - | 16 |
| 10 | NT female brood | adult (NT) | Thailand |  | TiLV (+) | - | 16 |
| 11 | NT male brood | adult (NT) | Thailand |  | TiLV (+) | - | 16 |
| 12 | NT juvenile 4 | juvenile (NT) | Thailand |  | TiLV (+) | - | 16 |
| 13 | RT F1 | fingerling (RT) | Thailand | 2015 | FnO (+) | - | 20 |
| 14 | RT F2 | fingerling (RT) | Thailand |  | FnO (+) | - | 20 |
| 15 | RT F3 | fingerling (RT) | Thailand |  | FnO (+) | - | 20 |
| 16 | RT F4 | fingerling (RT) | Thailand |  | FnO (+) | - | 20 |
| 17 | NT 1.2 | fingerling (NT) | Thailand | 2015 | TiLV (+) | - | 21 |
| 18 | NT 1.3 | fingerling (NT) | Thailand |  | TiLV (+) | - | 21 |
| 19 | NT 2.1 | fingerling (NT) | Thailand |  | TiLV (+) | - | 21 |
| 20 | NT 2.3 | fingerling (NT) | Thailand |  | TiLV (+) | - | 21 |
| 21 | NT 3.1 | fingerling (NT) | Thailand |  | TiLV (+) | - | 21 |
| 22 | NT 3.3 | fingerling (NT) | Thailand |  | TiLV (+) | - | 21 |
| 23 | NT Gh 21 | juvenile (NT) | West Africa | 2019 | ISKNV (++) | ISKNV* | - |
| 24 | NT Gh 22 | juvenile (NT) | West Africa |  | ISKNV (++) | ISKNV* | - |
| 25 | *r2_NT Gh 23* | juvenile (NT) | West Africa |  | ISKNV (++) | ISKNV* | - |
| 26 | *r2_NT Gh 24* | adult (NT) | West Africa |  | ISKNV (++) | ISKNV* | - |
| 27 | *r2_NT Gh 25* | juvenile (NT) | West Africa |  | ISKNV (++) | ISKNV* | - |
| 28 | RT EX 01 | fry (RT) | Thailand | 2019 | FnO (++) | FnO# | 22 |
| 29 | RT EX02 | fry (RT) | Thailand |  | FnO (++) | FnO# | 22 |
| 30 | *r2_RT EX03* | fry (RT) | Thailand |  | FnO (++) | FnO# | 22 |
| 31 | *r2_RT FM1a* | adult (RT) | Thailand |  | FnO (++) | FnO# | 22 |
| 32 | *r2_RT M1* | adult (RT) | Thailand |  | FnO (++) | FnO# | 22 |
| 33 | *r2_RT FM1b* | adult (RT) | Thailand |  | FnO (++) | FnO# | 22 |
| 34 | NT PV F8 | adult (NT) | Thailand | 2020 | Not done | * | - |
| 35 | NT PV F9 | adult (NT) | Thailand |  | Not done | # | - |
| 36 | NT PV F10 | adult (NT) | Thailand |  | Not done | # | - |
| 37 | *r3_NT Group B* | fry (NT) | Thailand | 2019 | SAG (++) | SAG | - |
| 38 | *r3_NT Zone A* | fry (NT) | Thailand |  | SAG (++) | SAG* | - |
| 39 | *r3_NT Zone B* | fry (NT) | Thailand |  | SAG (++) | SAG | - |
| 40 | *r1_TiLV_1* | fingerling (RT) | Thailand | 2018-07 | TiLV | TiLV | 23 |
| 41 | *r1_ISKNV_1* | fry (AS) | Thailand | 2018-11 | ISKNV | ISKNV | 24 |
| 42 | *r1_FNO_1* | fingerling (RT) | Thailand | 2015 | FnO | FnO | 20 |
| 43 | *r1_SAG_1* | fry (NT) | Thailand | 2019-07 | SAG | SAG |  |
| 44 | *r3_TiLV50* | fingerling (RT) | Thailand | 2019-07 | TiLV | TiLV | TiLV PCR amplicon of 100 ng |
| 45 | *r3_TiLV30* | fingerling (RT) | Thailand | 2019-07 | TiLV | TiLV | TiLV PCR amplicon of 75 ng |
| 46 | *r3_TiLV15* | fingerling (RT) | Thailand | 2019-07 | TiLV | TiLV | TiLV PCR amplicon of 50 ng |
| 47 | *r3_TiLV05* | fingerling (RT) | Thailand | 2019-07 | TiLV | TiLV | TiLV PCR amplicon of 25 ng |
| 48 | *r3_SAG50* | fry (NT) | Thailand | 2019-07 | SAG | SAG | SAG PCR amplicon of 100 ng |
| 49 | *r3_SAG30* | fry (NT) | Thailand | 2019-07 | SAG | SAG | SAG PCR amplicon of 75 ng |
| 50 | *r3_SAG15* | fry (NT) | Thailand | 2019-07 | SAG | SAG | SAG PCR amplicon of 50 ng |
| 51 | *r3_SAG05* | fry (NT) | Thailand | 2019-07 | SAG | SAG | SAG PCR amplicon of 25 ng |
| 52 | *r1_4PAT_1* | N/A | Thailand | N/A | 4PAT | 4PAT | PCR amplicons 4 PAT (combined1) |
| 53 | *r1_4PAT_DIL_1* | N/A | Thailand | N/A | 4PAT | 4PAT | PCR amplicons 4 PAT (combined2) |
| 54 | *r3_4PAT_DIL_3* | N/A | Thailand | N/A | 4PAT | 4PAT | PCR amplicons 4 PAT (combined3) |

Notes and abbreviations: sPCR, singleplex PCR; mPCR, multiplex PCR; NT, Nile tilapia; RT, red tilapia; AS, Asian sea bass; -, negative test; ++, high pathogen load; +, low pathogen load; \*, suspected dual infections with other pathogen(s); #, probably non-specific products; TiLV, tilapia lake virus; ISKNV, infectious spleen and kidney necrosis virus; FnO, *Francisella noatunensis* subsp. *orientalis*; SAG, *Streptococcus agalactiae*; 4PAT: mixture of PCR amplicons 4 pathogens (TiLV, ISKNV, FnO, SAG). Samples sequenced are italicized and highlighted in grey, and the designation "r1" to "r3" indicates the run number in Nanopore.

**Fig. 1.**
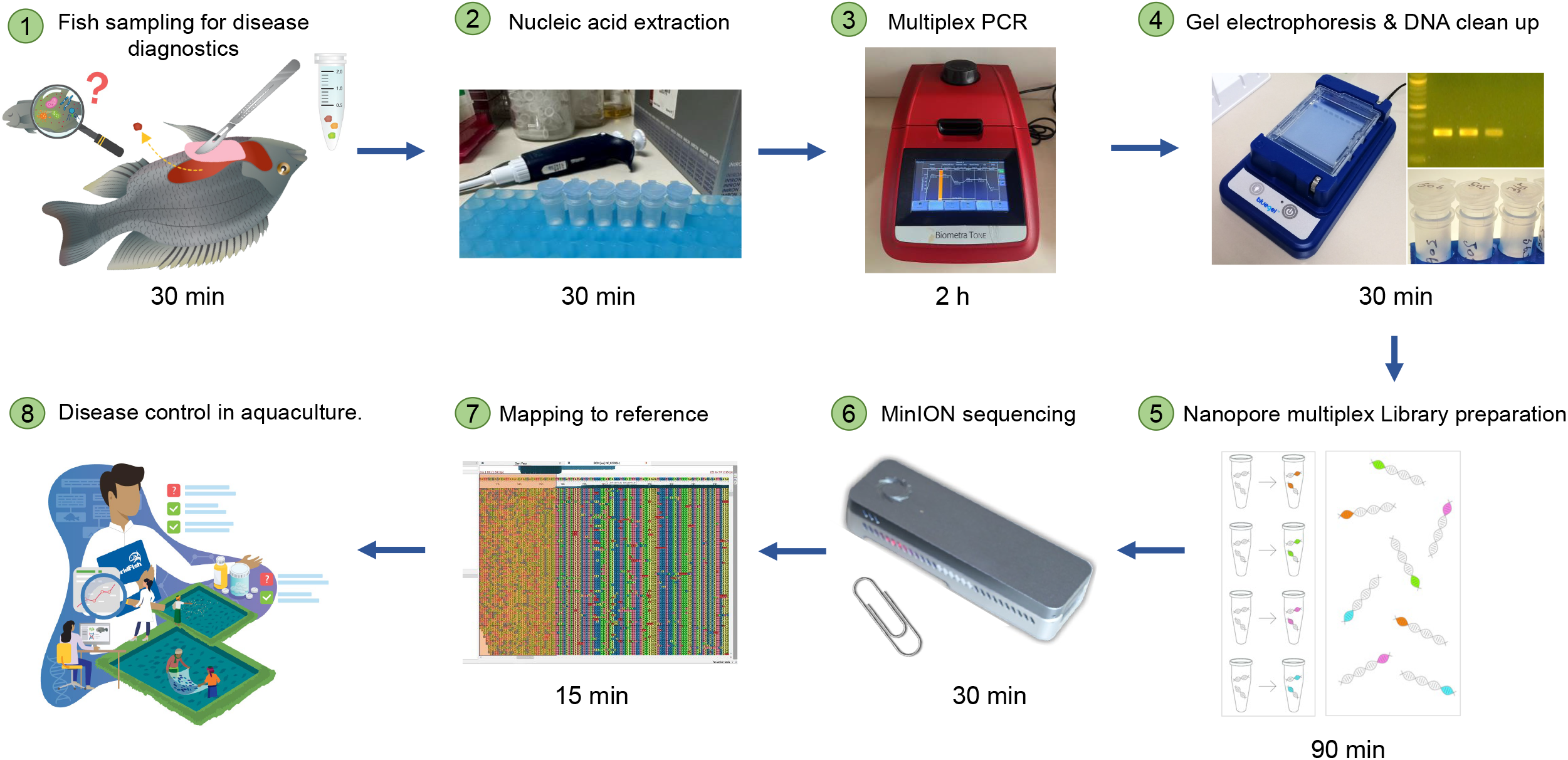

### Primers used in this study

We obtained the primers for the four selected target pathogens from Bio Basic (Canada), and the primer sequences are summarized in Supplemental Table 1. Primers for TiLV, ISKNV, *F. orientalis*, and *S. agalactiae* were reported in previous studies (Yang et al., 2013; Dong et al., 2016; Paria et al., 2016; Leigh et al., 2018; Kawato et al., 2021; Taengphu et al., 2022). The expected amplicon sizes were 137, 190, 203, and 351 bp, respectively. The specificity of each primer pair was assessed *in silico* using the Primer-BLAST program (https://www.ncbi.nlm.nih.gov/tools/primer-blast/).

### Plasmid positive controls and the analytical sensitivity assay

All plasmid-positive controls except for *S. agalactiae* (constructed in this study) were obtained from our previous studies (Supplemental Table 2). Positive plasmid control containing *S. agalactiae groEL* partial gene was obtained by cloning a 351 bp-*groEL* amplified fragment purified before being ligated into pGEM-T easy vector (Promega). A recombinant plasmid was subjected to DNA sequencing (Macrogen). The copy number of each plasmid was calculated based on its size in base pair (bp) and amount in nanogram (ng) using a web tool at http://cels.uri.edu/gsc/cndna.html. A combination of four recombinant plasmids was mixed for the multiplex PCR sensitivity assay. This mixture was then subjected to 10-fold serial dilution, resulting in a 1 to 10^6^ copies/μl range. Subsequently, 4 μl of the diluted series was used in the multiplex PCR reaction. To simulate a clinical sample from a fish, each PCR reaction was spiked with 50 ng of RNA extracted from a healthy tilapia.

### Multiplex PCR condition optimization and the detection of clinical samples

Previous studies claimed that qPCR reagents usually contain PCR additives and enhancers that can improve amplification efficiency (Karunanathie et al., 2022). To optimize our multiplex PCR, we used KAPA SYBR FAST One-Step qRT-PCR master mix (Roche), known to contain such additives. Each reaction was prepared in a 25 μl volume, comprising 1X master mix, 4 μl of template, and 80-240 nM of each primer pair (Supplemental Table 3). We determined the optimal annealing temperature (Ta) using gradient PCR with Ta ranging between 55-65 °C. To identify the best combination of PCR components, we added ammonium sulfate, BSA, dNTPs, and MgCl2 in different proportions. The final cycling conditions comprised a reverse transcription step at 42 °C for 5 min, followed by inactivation at 95 °C for 3 min, and 40 cycles of 95°C for 10 s and 60°C for 30 sec, with a final extension step at 60 °C for 5 min (Supplemental Table 3). We then analyzed 10 μl of each product by electrophoresis on a 3.5% agarose gel stained with ethidium bromide. The newly optimized multiplex PCR assay was subsequently used to detect the presence of TiLV, ISKNV, *F. orientalis*, and *S. agalactiae* in archival clinical samples (Table 1). Most of these samples were previously tested for a single pathogen using PCR or qPCR assays.

### Quantification of pathogens by qPCR assays

A total of 15 clinical samples, whose mPCR products underwent Nanopore sequencing, were analyzed to determine the presence and quantity of each pathogen. The established protocols (Leigh et al., 2018; Kawato et al., 2021; Taengphu et al., 2022) were followed to detect TiLV, ISKNV, and SAG, as described in Supplemental Table 4. For FnO qPCR, the same primers used in mPCR were used in this study’s SYBR Green-based qPCR assay.

### Nanopore sequencing

The present study employed amplicons generated from both single and multiplex PCR reactions (Table 1, samples highlighted in grey) as templates for library preparation using the ligation sequencing kit (SQK-LSK109) and the native barcoding expansion 1-12 kit (EXP-NBD104) according to the standard protocols of Oxford Nanopore Technologies (ONT) adapted for the Flongle flow cell. Three sequencing runs (r1-r3) were conducted, with 250 ng of PCR product per sample and a unique native barcode (BC) assigned to each sample. Run 1 (r1) involved sequencing amplicons obtained from individual PCR reactions and a combination of these amplicons (Table 1). Run 2 (r2) focused on sequencing multiplex PCR (mPCR) products derived from clinical samples. Run 3 (r3) employed mPCR products from the remaining clinical samples. Additionally, different concentrations of single PCR products were included in this run. The library of pooled barcoded samples was subjected to a Short Fragment Buffer (SFB) wash before the final elution step of the protocol. The DNA concentration was quantified at every step using the Qubit assay. Subsequently, the prepared library was loaded onto a Flongle flow cell (FLO-FLG106), following the Nanopore standard protocol, and each Flongle flow cell was fitted to a Flongle adapter (FLGIntSP) for MinION.

### Reads filtering and read abundance calculation

Basecalled reads in fastq format were primer-trimmed with cutadapt v4.3 (Martin, 2011), and reads that have been trimmed with length ranging from 75-400 bp will be retained for the subsequent analysis, leaving out overly short or long reads without intact primer sequence on both ends. The filtered reads were aligned to the four pathogen gene segments using Minimap2 v2.17 (Li, 2018). Reads that aligned uniquely (only a single hit reported) with more than 80% coverage to the target region were used for abundance calculation and variant calling (consensus generation). Calculation of raw, trimmed, and aligned read statistics used seqkit v2.20 (Shen et al., 2016). Reads failing to align at this stringent level were extracted and re-aligned using the default Minimap2 setting, followed by a less stringent blastN alignment. The host genome was included in the reference sequence to assess the fraction of reads mapping to the host.

### Generation of consensus and variant sequences

Only reads with unique alignments were used as the template for Minimap2 and medaka variant calling based on the ARTIC pipeline (“Core Pipeline -artic pipeline”; “sars-cov-2-ont-artic-variant-calling/COVID-19-ARTIC-ONT”). The reads were mapped to the reference gene segments of four pathogens, followed by variant calling using the medaka variant model r941_min_g507. Variant filtering was then performed based on mapping quality and read depth. The number of ambiguous bases in the consensus sequences was calculated using QUAST v5 (Gurevich et al., 2013). Only sequences without any ambiguous base were selected for subsequent variant sequence generation. Variants were generated using cd-hit v.4.8.1 (Huang et al., 2010), whereby sequences with 100% nucleotide identity and exact sequence length were clustered into the same genotype. The variant sequences were compared against the NCBI Blast nt database (accessed on 25 April 2023) to identify the top 5 BLAST hits for each sequence.

### Code availability

The Linux scripts used to generate initial FastQ files, assembled amplicons (public and from this study) are publicly available in the Zenodo.org dataset https://doi.org/10.5281/zenodo.7866295.

## Results

### Primer testing by single PCR

Single PCR reactions were performed using each primer pair with its respective target template to validate the selected primer pairs for each pathogen. The amplification of the target regions was confirmed by amplicons of the expected sizes, as illustrated in Supplemental Figure 1. To simulate the expected products from multiplex PCR (mPCR), an equal amount of each individual PCR product was loaded into lane C of the Supplemental Figure 1. Gel electrophoresis analysis revealed distinct and separable bands, demonstrating the potential for further utilization in the multiplex PCR analysis. Note that the products generated from this step were subsequently employed in Nanopore sequence run 1 (r1).

### Multiplex PCR detection of four pathogens among clinical samples

We have successfully optimized a multiplex PCR assay to simultaneously detect four important tilapia pathogens, TiLV, ISKNV, *F. orientalis*, and *S. agalactiae*, in a single reaction. The detection sensitivity of the new assay in the presence of spiked host RNA was 10^3^ copies/reaction for TiLV, ISKNV, and *S. agalactiae*. In the presence of 10^4^ copies of each template, the assay could detect *F. orientalis* and three other pathogens (Supplemental Figure 2). The assay could detect each pathogen efficiently in tilapia clinical samples, as confirmed by distinct PCR bands (Figure 2). The assay was able to detect TiLV in heavily infected samples (3-7) both on the gel and by nanopore but failed to detect it on the gel for samples (8-12) with low levels of TiLV infection. Similarly, the highest number of reads for ISKNV were found in sample #25 (twice more than in sample #27), yet it had a much fainter band on the gel (Table 1). There is a good correlation between read numbers and bands intensity for *F. orientalis* in four samples (#30-33) but only a weak band for *S. agalactiae* in one sample (#39) compared to number of amplicons (Figure 2, 3, Supplemental Table 5). In some samples, we detected dual infections with more than one pathogen, as evidenced by the presence of multiple bands, including a possible co-infection with *F. orientalis* and TiLV in samples 3-7. However, differentiation of *F. orientalis* and ISKNV was challenging as their respective bands have similar sizes (Figure 2).

**Fig. 2.**
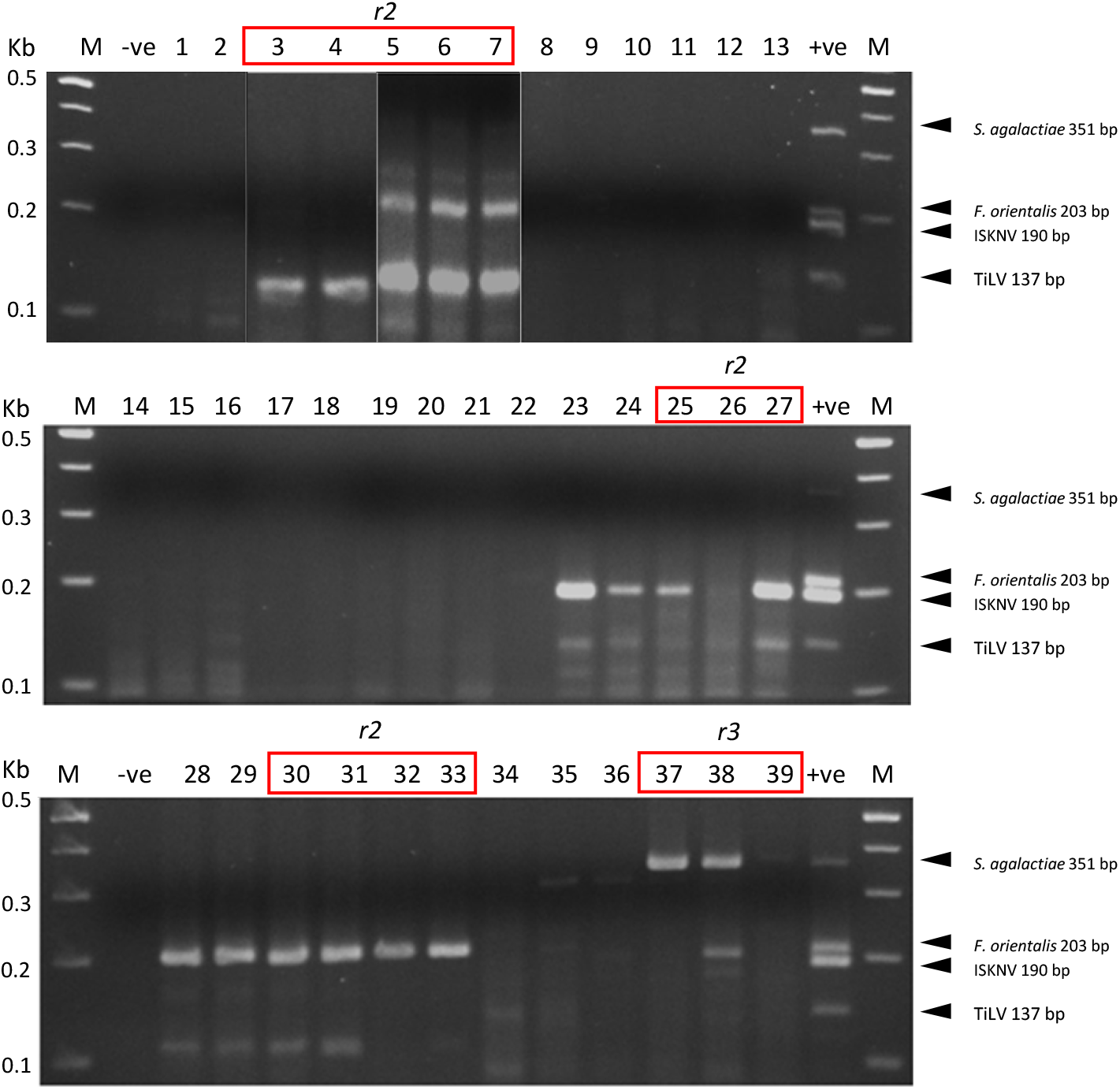

**Fig. 3.**
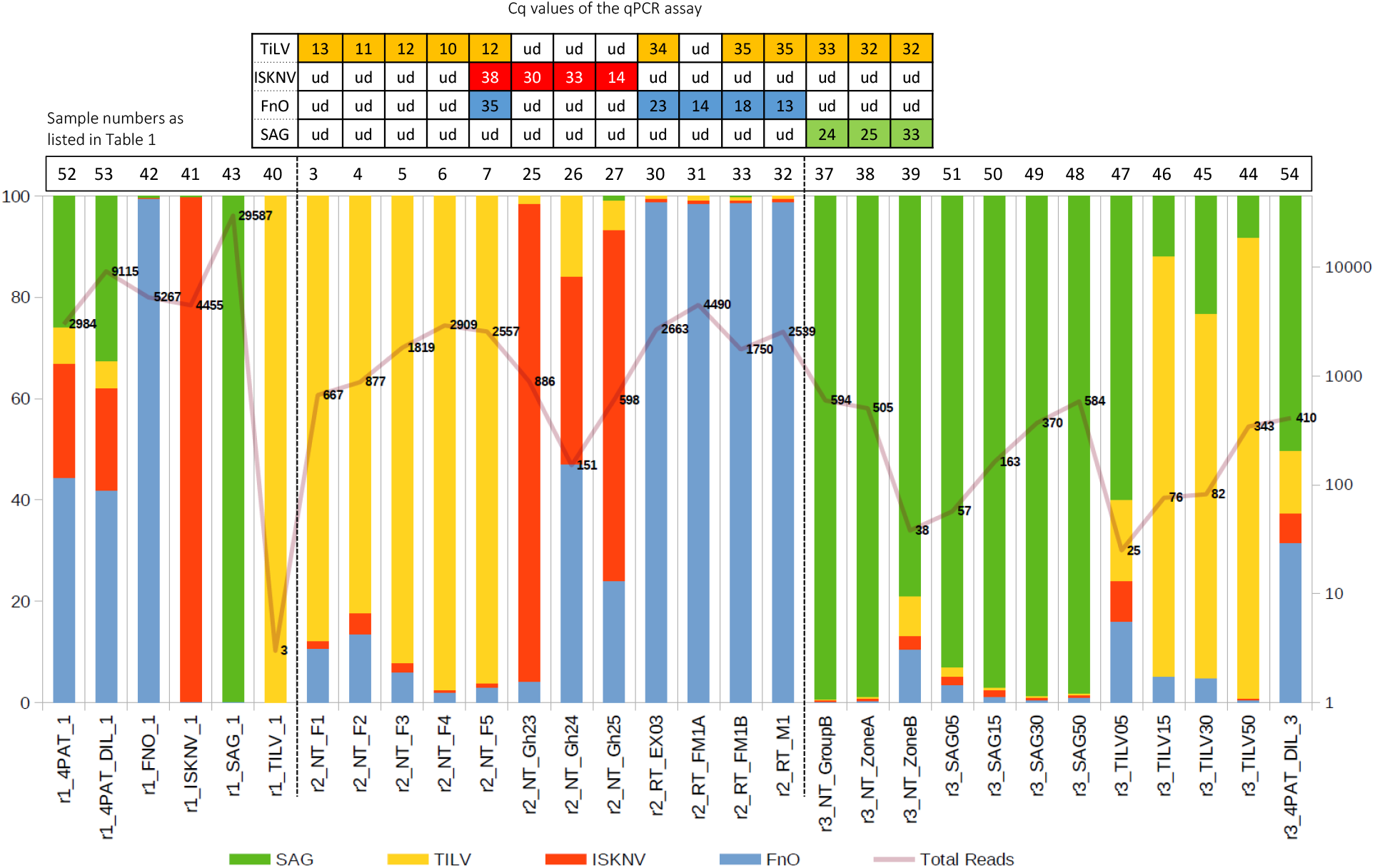

### Successful recovery of pathogen sequence variants from infected samples

A total of 246,756 reads were successfully demultiplexed from three separate Flongle runs (r1-r3), each containing different sample types (Table 1, Supplemental Table 5). Following primer and length filtering, a substantial reduction in read count was observed, resulting in an average of 64% of reads being removed (range: 46%-87%) (Supplemental Table 5). An additional 10% of reads were removed after alignment filtering. After these filtering steps, 75,203 reads remained, providing an average of 2,500 filtered reads per sample for subsequent read abundance and consensus sequence generation (Supplemental Table 5).

Given the small sampling size and possibly highly conserved nature of some of the pathogen gene segments, one sequence variant was generated per pathogen gene segment except for the TiLV gene segment, whereby two sequence variants were recovered (Table 2). All recovered sequence variants exhibited 100% nucleotide identity to at least one publicly available sequence in the NCBI database. In addition, samples derived from the single PCR product inputs and their combined mixture were clustered in their respective sequence variants. In the case of the TiLV amplicons, we observed two unique sequence variants (Var 0 and Var 1) for TiLV gene segment 9, one of which displayed 100% nucleotide identity to the gene region of a relatively divergent TiLV strain from Vietnam (Table 2). This sequence variant was detected in samples r2_NT_F1 to r2_NT_F5, collected from the same sampling site simultaneously (Table 2).

**Table 2.** List of variants for the pathogen gene region, along with the inferred variants and their top 5 NCBI BLAST hits. Pat: Pathogen; Var, Variants; %ID, % nucleotide identity. The samples were grouped based on the sequencing run from which they were generated.

| Pat | Samples | Var | Top 5 BLAST hits | %ID | Accession |
| --- | --- | --- | --- | --- | --- |
| FnO | r1_4PAT_1,r1_4PAT_DIL_1,r1_FNO_1,r1_SAG_1<br>r2_NT_F1,r2_NT_F2,r2_NT_F3,r2_NT_F4,r2_NT_F5,r2_NT_Gh23,r2_NT_Gh24,r2_NT_Gh25,r2_RT_EX03,r2_RT_FM1A,r2_RT_FM1B,r2_RT_M1<br>r3_4PAT_DIL_3 | 0 | *Francisella noatunensis* subsp. *orientalis* strain LPM2-AR2019 | 100 | MN385384.1 |
|  |  |  | *Francisella noatunensis* subsp. *orientalis* strain FO371 | 100 | CP022953.1 |
|  |  |  | *Francisella noatunensis* subsp. *orientalis* strain FNO364 | 100 | CP022952.1 |
|  |  |  | *Francisella noatunensis* subsp. *orientalis* strain FNO222 | 100 | CP022951.1 |
|  |  |  | *Francisella noatunensis* subsp. *orientalis* strain FNO215 | 100 | CP022950.1 |
| ISKNV | r1_4PAT_1,r1_4PAT_DIL_1,r1_ISKNV_1<br>r2_NT_F2,r2_NT_Gh23,r2_NT_Gh24,r2_NT_Gh25 | 0 | ISKNV isolate ISKNV/10 | 100 | MT178422.1 |
|  |  |  | ISKNV isolate ISKNV/48 | 100 | MT178418.1 |
|  |  |  | Angelfish iridovirus AFIV-16 | 100 | MK689685.1 |
|  |  |  | ISKNV isolate M6 | 100 | MK084827.1 |
|  |  |  | ISKNV isolate SB04 | 100 | KY440040.1 |
| SAG | r1_SAG_1,r1_4PAT_1,r1_4PAT_DIL_1,r1_SAG_1<br>r3_4PAT_DIL_3,r3_NT_GroupB,r3_NT_ZoneA,r3_NT_ZoneB,r3_SAG05,r3_SAG15,r3_SAG30,r3_SAG50 | 0 | *Streptococcus agalactiae* strain 01173 | 100 | CP053027.1 |
|  |  |  | *Streptococcus agalactiae* strain Sag153 | 100 | CP036376.1 |
|  |  |  | *Streptococcus agalactiae* strain ZQ0910 | 100 | CP049938.1 |
|  |  |  | *Streptococcus agalactiae* strain NJ1606 | 100 | CP026084.1 |
|  |  |  | *Streptococcus agalactiae* strain BSE009 | 100 | CP020387.1 |
| TiLV | r1_4PAT_1,r1_4PAT_DIL_1<br>r3_4PAT_DIL_3,r3_TiLV15,r3_TiLV30,r3_TiLV50 | 0 | TiLV isolate WVL19054 segment 9 | 100 | MN193531.1 |
|  |  |  | TiLV isolate WVL19031-01A segment 9 | 100 | MN193521.1 |
|  |  |  | TiLV isolate EC-2012 segment 9 | 100 | MK392380.1 |
|  |  |  | TiLV isolate Til-4-2011 segment 9 | 100 | KU751822.1 |
|  |  |  | TiLV AD-2016 Contig 20 | 100 | KU552140.1 |
|  | r2_NT_F1,r2_NT_F2,r2_NT_F3,r2_NT_F4,r2_NT_F5 | 1 | TiLV isolate RIA2-VN-2019 segment 9 | 100 | ON376590.1 |
|  |  |  | TiLV isolate WVL19054 segment 9 | 98.91 | MN193531.1 |
|  |  |  | TiLV isolate WVL19031-01A segment 9 | 98.91 | MN193521.1 |
|  |  |  | TiLV strain EC-2012 segment 9 | 98.91 | MK392380.1 |
|  |  |  | TiLV isolate Til-4-2011 segment 9 | 98.91 | KU751822.1 |

Abbreviations: FnO, *Francisella noatunensis* subsp. *orientalis*; ISKNV, infectious spleen and kidney necrosis virus; SAG, *Streptococcus agalactiae*; TiLV, tilapia lake virus

### Effect of template input on data distribution and pathogen detection specificity on the Nanopore platform

In the first run (r1), using well-defined pathogen nucleic acids as templates for the detection assay, confident alignments were observed for each sample to their expected pathogen gene segments, except for sample r1_TILV_1 which consisted purely of TiLV amplicons and did not sequence well enough to enable subsequent variant calling (Figure 3 and Table 2). For sample 4PAT, which consisted of a combination of products from 4 pathogens, the read distribution was broadly similar, as was the case for sample 4PAT_DIL, which used a ten-fold diluted version of the template (Figure 3). In the second run (r2), actual clinical samples with PCR-verified infections were used, and over 50% and up to 99% of reads were primarily mapped to the primary suspected pathogen gene fragments, with the remaining reads mapping to the other three pathogens. Interestingly, in the third run (r3), a significant percentage (>5%) of reads aligned to other pathogens were observed among samples spiked with different amounts of single amplified products (Figure 3). For example, among the TiLV samples (TILV05, 15, 30, 50), the lowest amount of PCR product template (TILV05) resulted in a low number of TiLV reads and a higher percentage of reads belonging to non-TiLV pathogens. A similar trend was observed for pure SAG samples, with the most diluted SAG sample (SAG05) having the highest percentage of non-SAG reads. However, in contrast to the pure synthetic TiLV samples, the percentage of non-specific reads decreased to less than 2% in the SAG50 sample with the highest read count, while it remained around 10% in the TILV50 sample.

### Reads that failed stringent alignment to the pathogen gene panel were host-derived or partial pathogen sequences with lower nucleotide identity

Upon investigation of reads that failed to align, a significant proportion was found to map not only to the four pathogen gene segments but also to the host genome when a more lenient alignment configuration was employed (Figure 4). Samples from run 2 (r2), predominantly derived from fish tissues, exhibited a higher percentage of reads mapping to the host genome than other samples. As expected, given the use of input PCR products as the template for all samples in run 1 (r1), no reads were found to align to the host genome when they were aligned with Minimap2 (default setting). On the contrary, a small portion of reads belonging to the host genome was found among different concentrations of single PCR products from run 3 (r3) (Figure 4 and Supplemental Table 5), which also consists of samples derived from tilapia organs (Table 1).

**Fig. 4.**
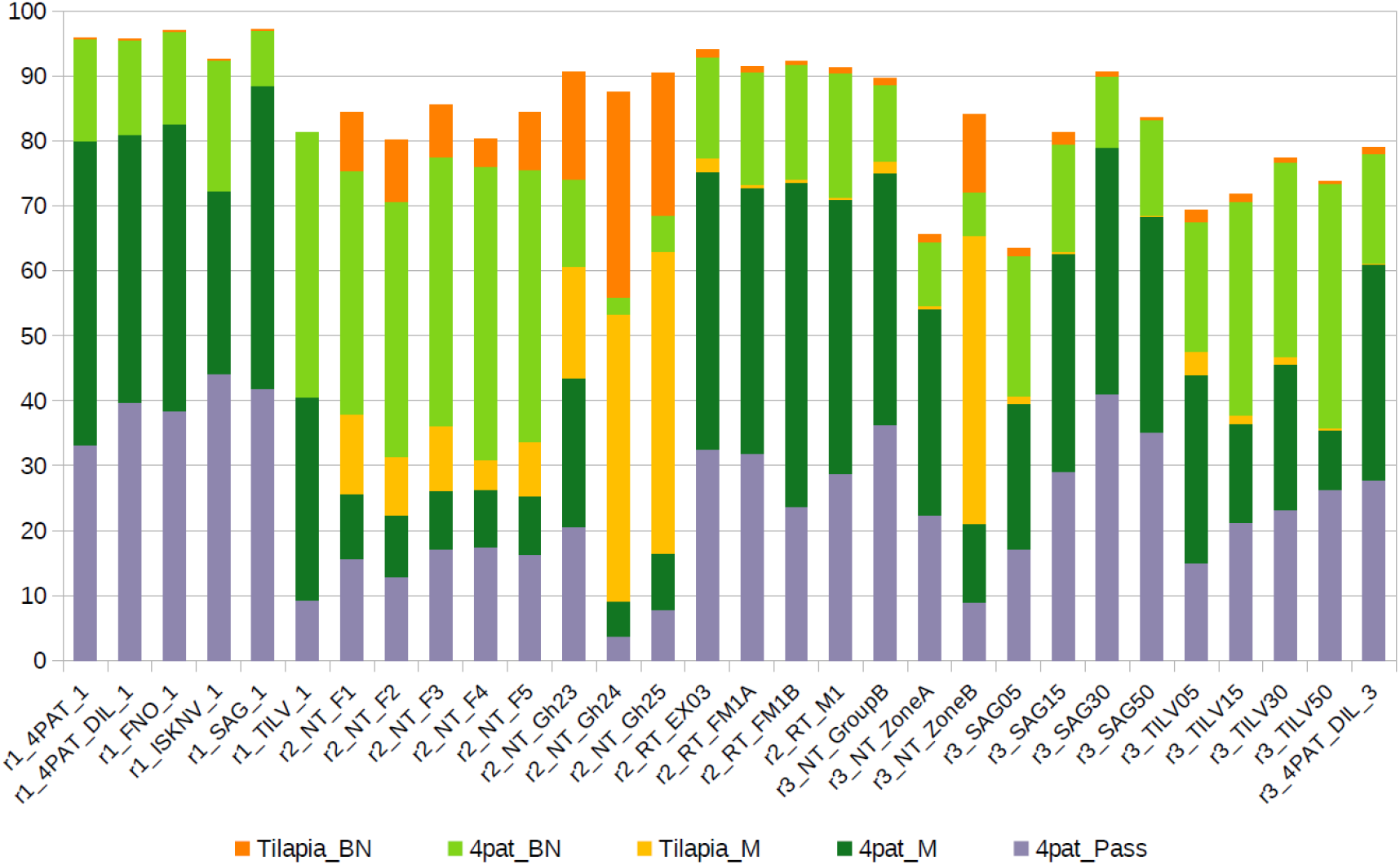

## Discussion

The intensification of tilapia farming has led to a surge in emerging and re-emerging infectious diseases, resulting in substantial losses in the aquaculture industry (Machimbirike et al., 2019; Haenen et al., 2023; Shinn et al., 2023). These diseases continue to evolve and spread under intense selective pressures, particularly in regions with high-density tilapia farming. To address this issue, we built upon our previous study (Delamare-Deboutteville et al., 2021), and improved the detection and genotyping of multiple pathogens by switching to multiplex PCR coupled with Nanopore sequencing. Our updated approach enabled the simultaneous and sequence-based detection and verification of four major tilapia pathogens: ISKNV, TiLV, *S. agalactiae*, and *F. orientalis*.

The requirement for simultaneous diagnosis of multiple pathogens is particularly relevant to high mortalities observed in larger grow-out tilapia. These mortalities are likely exacerbated by co-infections, as indicated by (Ramírez-Paredes et al., 2021) previous research findings, which revealed the active co-infection of ISKNV-positive fish with Streptococcus agalactiae and other bacterial pathogens.

The discrepancies between gel-based interpretation of multiplex PCR products and qPCR or direct amplicon sequencing approaches are not surprising, given the subjective nature of gel visualization and the labor-intensive process of running multiple samples. The accuracy of pathogen detection can be significantly impacted by faint or similar-sized bands, making it challenging to standardize results. To streamline the multiplex PCR approach and increase throughput, the gel visualization step can be eliminated in favor of direct Nanopore sequencing. With high-degree sample-based multiplexing, it is now possible to perform high throughput sequence-based detection of multiple tilapia pathogens using Nanopore sequencing. The utilization of this application is paramount for the aquaculture industry, given that disease outbreaks are frequently linked with multiple infections (AbdellLatif & Khafaga, 2020; Huang et al., 2020; Liu et al., 2020; Basri et al., 2020). Furthermore, as aquaculture farms are often situated in remote locations far from diagnostic reference laboratories, the ability to mobilize diagnostic testing with rapid results, high accuracy, and less complicated equipment near the farm is preferred.

Although our approach is scalable, there are limitations due to the old version of MinKNOW used during the study. Short amplicons are sequenced less efficiently and have lower read accuracy. We addressed this issue using a stringent read filtering and alignment approach, which led to a reduced read recovery in the final abundance calculation. Some synthetic samples analyzed contained reads from other pathogens, which was unexpected and suggests cross-ligation of native barcodes during library preparation. The availability of unbound adapters is higher among samples with low DNA input, leading to higher crosstalk levels and reads from other samples. New improvements in the current Nanopore technology and library preparation protocols (as of 14 September 2024) will mitigate these issues. This includes (1) Switching to a PCR-barcoding kit to only enrich for amplicon with Nanopore partial adapter, (2) the use of a new Q20+ LSK114 kit with improved read accuracy, and (3) the use of high accuracy Dorado base-calling model (https://github.com/nanoporetech/dorado).

By enforcing a stringent alignment setting, the reads aligning to the pathogen gene segment are directly suitable for subsequent reference-based variant calling, producing highly accurate variants useful for biological interpretation. This is exemplified by the consistent recovery of a TiLV segment 9 variant previously only found in a Vietnamese TiLV strain in all Thai tilapia samples from the same sampling batch. Considering the proximity of Thailand and Vietnam, this may suggest a possible spillover event, highlighting the need for more thorough sampling and sequencing to elucidate the degree of TiLV diversity in both regions.

For field diagnostics, some of the equipment used in the present study can be replaced with portable items that are small enough to fit in a bag for use in remote settings. An example of the field application of this diagnostic workflow is the Lab-in-a-backpack concept developed by WorldFish (WorldFish, 2020; Huso et al., 2020; Cagua et al., 2021; The University of Queensland, 2021; Chadag et al., 2021; Barnes et al., 2021). It includes all the necessary sampling, extraction, and sequencing kits, pipettes, and consumables, as well as the use of the miniPCR thermocycler and blueGel electrophoretic system from miniPCR bio™, a minicentrifuge, a magnetic rack, a fluorometer, a Nanopore MinION sequencer, and flow cells for sequencing. Additionally, a laptop with minimum requirements to run the MinKNOW software is required. Similar PCR-sequencing approaches using portable equipment have been employed for the rapid identification of a diverse range of biological specimens, including plants, insects, reptiles (Pomerantz et al., 2022), and viruses such as COVID-19, Ebola, and Zika (González-González et al., 2019, 2020).

Improvements in the remote and low-resource applications of the current methodology will broaden its use for capacity building and implementations in low-income countries, where high mortalities due to unknown causes are prevalent in tilapia farming. Accurate and rapid genomic detection of fish pathogens through our amplicon-based approach will allow health professionals to promptly provide advice and take necessary actions for producers. Moreover, it can facilitate informed decisions regarding additional investigations, such as whole-genome sequencing (WGS), on isolates stored in biobanks. The sequence data obtained from WGS offer crucial epidemiological insights, which can be utilized to develop customized multivalent autogenous vaccines using local pathogens (Barnes et al., 2022). These vaccines can then be administered to broodstock and seeds before distribution among grow-out farmers for restocking purposes.

## Conclusions

In summary, our study presents an attractive approach for detecting and verifying four tilapia pathogens in clinically sick fish, which can also be applied to pathogens in other livestock, fish, or crustaceans if the genetic information of the pathogen is publicly available for primer design and reference mapping. Although there are limitations to the current pipeline version used at the time of this study, we are optimistic that the current improvements in Nanopore technology will further enhance the accuracy and scalability of our approach.

## Acknowledgements

We want to extend our utmost gratitude and profound recognition to the late Dr. Pattanapon Kayansamruaj for his valuable contributions during the early stages of conceiving this research endeavor.

## References

Abdel□Latif HMR, Khafaga AF. 2020. Natural co-infection of cultured Nile tilapia Oreochromis niloticus with Aeromonas hydrophila and Gyrodactylus cichlidarum experiencing high mortality during summer. Aquaculture Research 51:1880–1892. DOI: 10.1111/are.14538.

Alathari S, Chaput DL, Bolaños LM, Joseph A, Jackson VLN, Verner-Jeffreys D, Paley R, Tyler CR, Temperton B. 2023. A Multiplexed, Tiled PCR Method for Rapid Whole-Genome Sequencing of Infectious Spleen and Kidney Necrosis Virus (ISKNV) in Tilapia. Viruses 15:965. DOI: 10.3390/v15040965.

Barnes A, Das S, Rudenko O, Wilkinson S, Cagua F, Delamare-Deboutteville J. 2021. Diagnosis in a fish farmer’s backpack. In: Food & Nutrition Security – The Biosecurity, Health, Trade Nexus. https://www.crawfordfund.org/events/other-events/2021-conference/speakers-chairs/dr-andrew-barnes/. Crawford Fund,.

Barnes AC, Silayeva O, Landos M, Dong HT, Lusiastuti A, Phuoc LH, Delamare-Deboutteville J. 2022. Autogenous vaccination in aquaculture: A locally enabled solution towards reduction of the global antimicrobial resistance problem. Reviews in Aquaculture 14:907–918. DOI: 10.1111/raq.12633.

Basri L, Nor RM, Salleh A, Md Yasin IS, Saad MZ, Abd Rahaman NY, Barkham T, Amal MNA. 2020. Co-Infections of Tilapia Lake Virus, Aeromonas hydrophila and Streptococcus agalactiae in Farmed Red Hybrid Tilapia. Animals 10:2141. DOI: 10.3390/ani10112141.

Cagua F, Wilkinson S, Barnes A, Delamare-Deboutteville J. 2021. Lab in a Backpack: Rapid and accurate diagnosis for the control and prevention of diseases in aquatic animals. Available at https://labinabackpack.com/ (accessed June 15, 2023).

Chadag V, Delamare-Deboutteville J, Beveridge M, Marwaha N. 2021. Reducing disease risks in fish through better detection, management and prevention. WorldFish. https://worldfishcenter.org/publication/reducing-disease-risks-fish-through-better-detection-management-and-prevention.

Core Pipeline - artic pipeline. Available at https://artic.readthedocs.io/en/latest/minion/ (accessed June 15, 2023).

Debnath SC, McMurtrie J, Temperton B, Delamare-Deboutteville J, Mohan CV, Tyler CR. 2023. Tilapia aquaculture, emerging diseases, and the roles of the skin microbiomes in health and disease. Aquaculture International. DOI: 10.1007/s10499-023-01117-4.

Delamare-Deboutteville J, Taengphu S, Gan HM, Kayansamruaj P, Debnath PP, Barnes A, Wilkinson S, Kawasaki M, Vishnumurthy Mohan C, Senapin S, Dong HT. 2021. Rapid genotyping of tilapia lake virus (TiLV) using Nanopore sequencing. Journal of Fish Diseases 44:1491–1502. DOI: 10.1111/jfd.13467.

Dong HT, Chaijarasphong T, Barnes AC, Delamare-Deboutteville J, Lee PA, Senapin S, Mohan CV, Tang KFJ, McGladdery SE, Bondad-Reantaso MG. 2023. From the basics to emerging diagnostic technologies: What is on the horizon for tilapia disease diagnostics? Reviews in Aquaculture 15:186–212. DOI: 10.1111/raq.12734.

Dong HT, Gangnonngiw W, Phiwsaiya K, Charoensapsri W, Nguyen VV, Nilsen P, Pradeep PJ, Withyachumnarnkul B, Senapin S, Rodkhum C. 2016. Duplex PCR assay and in situ hybridization for detection of Francisella spp. and Francisella noatunensis subsp. orientalis in red tilapia. Diseases of Aquatic Organisms 120:39–47. DOI: 10.3354/dao03021.

FAO. 2020. The State of World Fisheries and Aquaculture 2020: Sustainability in action. Rome, Italy: FAO, Rome. DOI: 10.4060/ca9229en.

FAO. 2022. FishStatJ: universal software for fishery statistical time series: aquaculture production 1950–2020. FAO, Rome.

González-González E, Mendoza-Ramos JL, Pedroza SC, Cuellar-Monterrubio AA, Márquez-Ipiña AR, Lira-Serhan D, Santiago GT, Alvarez MM. 2019. Validation of use of the miniPCR thermocycler for Ebola and Zika virus detection. PLOS ONE 14:e0215642. DOI: 10.1371/journal.pone.0215642.

González-González E, Santiago GT, Lara-Mayorga IM, Martínez-Chapa SO, Alvarez MM. 2020. Portable and accurate diagnostics for COVID-19: Combined use of the miniPCR thermocycler and a well-plate reader for SARS-CoV-2 virus detection. PLOS ONE 15:e0237418. DOI: 10.1371/journal.pone.0237418.

Gurevich A, Saveliev V, Vyahhi N, Tesler G. 2013. QUAST: quality assessment tool for genome assemblies. Bioinformatics 29:1072–1075. DOI: 10.1093/bioinformatics/btt086.

Haenen OLM, Dong HT, Hoai TD, Crumlish M, Karunasagar I, Barkham T, Chen SL, Zadoks R, Kiermeier A, Wang B, Gamarro EG, Takeuchi M, Azmai MNA, Fouz B, Pakingking Jr. R, Wei ZW, Bondad-Reantaso MG. 2023. Bacterial diseases of tilapia, their zoonotic potential and risk of antimicrobial resistance. Reviews in Aquaculture 15:154–185. DOI: 10.1111/raq.12743.

Huang Y, Cai S, Jian J, Liu G, Xu L. 2020. Co-infection of infectious spleen and kidney necrosis virus and Francisella sp. in farmed pearl gentian grouper (♀Epinephelus fuscoguttatus ×♂E. lanceolatus) in China - A case report. Aquaculture 526:735409. DOI: 10.1016/j.aquaculture.2020.735409.

Huang Y, Niu B, Gao Y, Fu L, Li W. 2010. CD-HIT Suite: a web server for clustering and comparing biological sequences. Bioinformatics (Oxford, England) 26:680–682. DOI: 10.1093/bioinformatics/btq003.

Huso D, Wilkinson S, Barnes A, Delamare-Deboutteville J. 2020. Lab-in-a-backpack: Rapid Genomic Detection to revolutionize control of disease outbreaks in fish farming. Available at https://fish.cgiar.org/impact/stories-of-change/lab-backpack-rapid-genomic-detection-revolutionize-control-disease (accessed June 11, 2021).

Karunanathie H, Kee PS, Ng SF, Kennedy MA, Chua EW. 2022. PCR enhancers: Types, mechanisms, and applications in long-range PCR. Biochimie 197:130–143. DOI: 10.1016/j.biochi.2022.02.009.

Kawasaki M, Delamare-Deboutteville J, Bowater RO, Walker MJ, Beatson S, Ben Zakour NL, Barnes AC. 2018. Microevolution of Streptococcus agalactiae ST-261 from Australia Indicates Dissemination via Imported Tilapia and Ongoing Adaptation to Marine Hosts or Environment. Applied and Environmental Microbiology 84. DOI: 10.1128/AEM.00859-18.

Kawato Y, Cummins DM, Valdeter S, Mohr PG, Ito T, Mizuno K, Kawakami H, Williams LM, Crane MSJ, Moody NJG. 2021. Development of New Real-time PCR Assays for Detecting Megalocytivirus Across Multiple Genotypes. Fish Pathology 56:177–186. DOI: 10.3147/jsfp.56.177.

Kralik P, Ricchi M. 2017. A Basic Guide to Real Time PCR in Microbial Diagnostics: Definitions, Parameters, and Everything. Frontiers in Microbiology 8.

Leigh WJ, Zadoks RN, Jaglarz A, Costa JZ, Foster G, Thompson KD. 2018. Evaluation of PCR primers targeting the groEL gene for the specific detection of Streptococcus agalactiae in the context of aquaculture. Journal of Applied Microbiology 125:666–674. DOI: 10.1111/jam.13925.

Li H. 2018. Minimap2: pairwise alignment for nucleotide sequences. Bioinformatics 34:3094–3100. DOI: 10.1093/bioinformatics/bty191.

Liamnimitr P, Thammatorn W, U-thoomporn S, Tattiyapong P, Surachetpong W. 2018. Non-lethal sampling for Tilapia Lake Virus detection by RT-qPCR and cell culture. Aquaculture 486:75–80. DOI: 10.1016/j.aquaculture.2017.12.015.

Liu X, Sun W, Zhang Y, Zhou Y, Xu J, Gao X, Zhang S, Zhang X. 2020. Impact of Aeromonas hydrophila and infectious spleen and kidney necrosis virus infections on susceptibility and host immune response in Chinese perch (Siniperca chuatsi). Fish & Shellfish Immunology 105:117–125. DOI: 10.1016/j.fsi.2020.07.012.

LópezLPorras A, Elizondo C, Chaves AJ, Camus AC, Griffin MJ, Kenelty K, Barum S, BarqueroLCalvo E, Soto E. 2019. Application of multiplex quantitative polymerase chain reaction methods to detect common bacterial fish pathogens in Nile tilapia, Oreochromis niloticus, hatcheries in Costa Rica. Journal of the World Aquaculture Society 50:645–658. DOI: 10.1111/jwas.12576.

Mabrok M, Elayaraja S, Chokmangmeepisarn P, Jaroenram W, Arunrut N, Kiatpathomchai W, Debnath PP, Delamare-Deboutteville J, Mohan CV, Fawzy A, Rodkhum C. 2021. Rapid visualization in the specific detection of Flavobacterium columnare, a causative agent of freshwater columnaris using a novel recombinase polymerase amplification (RPA) combined with lateral flow dipstick (LFD) assay. Aquaculture 531:735780. DOI: 10.1016/j.aquaculture.2020.735780.

Machimbirike VI, Jansen MD, Senapin S, Khunrae P, Rattanarojpong T, Dong HT. 2019. Viral infections in tilapines: More than just tilapia lake virus. Aquaculture 503:508–518. DOI: 10.1016/j.aquaculture.2019.01.036.

Martin M. 2011. Cutadapt removes adapter sequences from high-throughput sequencing reads. EMBnet.journal 17:10–12. DOI: 10.14806/ej.17.1.200.

Meemetta W, Domingos JA, Dong HT, Senapin S. 2020. Development of a SYBR Green quantitative PCR assay for detection of Lates calcarifer herpesvirus (LCHV) in farmed barramundi. Journal of Virological Methods 285:113920. DOI: 10.1016/j.jviromet.2020.113920.

Paria A, Dong J, Padinhate Purayil SB, Marappan M, Chaudhari A, Thirunavukkarasu AR, Purushothaman C, Kv R. 2016. Evaluation of candidate reference genes for quantitative expression studies in Asian seabass (Lates calcarifer) during ontogenesis and in tissues of healthy and infected fishes. Indian journal of experimental biology 54:597–605.

Pomerantz A, Sahlin K, Vasiljevic N, Seah A, Lim M, Humble E, Kennedy S, Krehenwinkel H, Winter S, Ogden R, Prost S. 2022. Rapid in situ identification of biological specimens via DNA amplicon sequencing using miniaturized laboratory equipment. Nature Protocols 17:1415–1443. DOI: 10.1038/s41596-022-00682-x.

Ramírez-Paredes JG, Paley RK, Hunt W, Feist SW, Stone DM, Field TR, Haydon DJ, Ziddah PA, Nkansa M, Guilder J, Gray J, Duodu S, Pecku EK, Awuni JA, Wallis TS, Verner-Jeffreys DW. 2021. First detection of infectious spleen and kidney necrosis virus (ISKNV) associated with massive mortalities in farmed tilapia in Africa. Transboundary and Emerging Diseases 68:1550–1563. DOI: 10.1111/tbed.13825.sars-cov-2-ont-artic-variant-calling/COVID-19-ARTIC-ONT. Available at https://workflowhub.eu/workflows/111 (accessed June 15, 2023).

Shen W, Le S, Li Y, Hu F. 2016. SeqKit: A Cross-Platform and Ultrafast Toolkit for FASTA/Q File Manipulation. PLOS ONE 11:e0163962. DOI: 10.1371/journal.pone.0163962.

Shinn AP, Avenant-Oldewage A, Bondad-Reantaso MG, Cruz-Laufer AJ, García-Vásquez A, Hernández-Orts JS, Kuchta R, Longshaw M, Metselaar M, Pariselle A, Pérez-Ponce de León G, Pradhan PK, Rubio-Godoy M, Sood N, Vanhove MPM, Deveney MR. 2023. A global review of problematic and pathogenic parasites of farmed tilapia. Reviews in Aquaculture 15:92–153. DOI: 10.1111/raq.12742.

Soto E, Bowles K, Fernandez D, Hawke JP. 2010. Development of a real-time PCR assay for identification and quantification of the fish pathogen Francisella noatunensis subsp. orientalis. Diseases of Aquatic Organisms 89:199–207. DOI: 10.3354/dao02204.

Taengphu S, Kayansamruaj P, Kawato Y, Delamare-Deboutteville J, Mohan CV, Dong HT, Senapin S. 2022. Concentration and quantification of Tilapia tilapinevirus from water using a simple iron flocculation coupled with probe-based RT-qPCR. PeerJ 10:e13157. DOI: 10.7717/peerj.13157.

The University of Queensland. 2021. Diagnosis in a fish farmer’s backpack. YouTube. https://www.youtube.com/watch?v=pUqmgQ-AuXM.

Waiyamitra P, Tattiyapong P, Sirikanchana K, Mongkolsuk S, Nicholson P, Surachetpong W. 2018. A TaqMan RT-qPCR assay for tilapia lake virus (TiLV) detection in tilapia. Aquaculture 497:184–188. DOI: 10.1016/j.aquaculture.2018.07.060.

Wang M, Lu M. 2016. Tilapia polyculture: a global review. Aquaculture Research 47:2363–2374. DOI: 10.1111/are.12708.

WorldFish. 2020. A New Concept for Rapid Genomic Detection of Fish Disease. YouTube. https://www.youtube.com/watch?v=iFxbodO6Fos.

Yang CG, Wang XL, Tian J, Liu W, Wu F, Jiang M, Wen H. 2013. Evaluation of reference genes for quantitative real-time RT-PCR analysis of gene expression in Nile tilapia (Oreochromis niloticus). Gene 527:183–192. DOI: 10.1016/j.gene.2013.06.013.

