## Supplementary figures and images for "Multiplex PCR with Nanopore Sequencing for Sequence-Based Detection of Four Tilapia Pathogens"

### Supplemental Figure 1

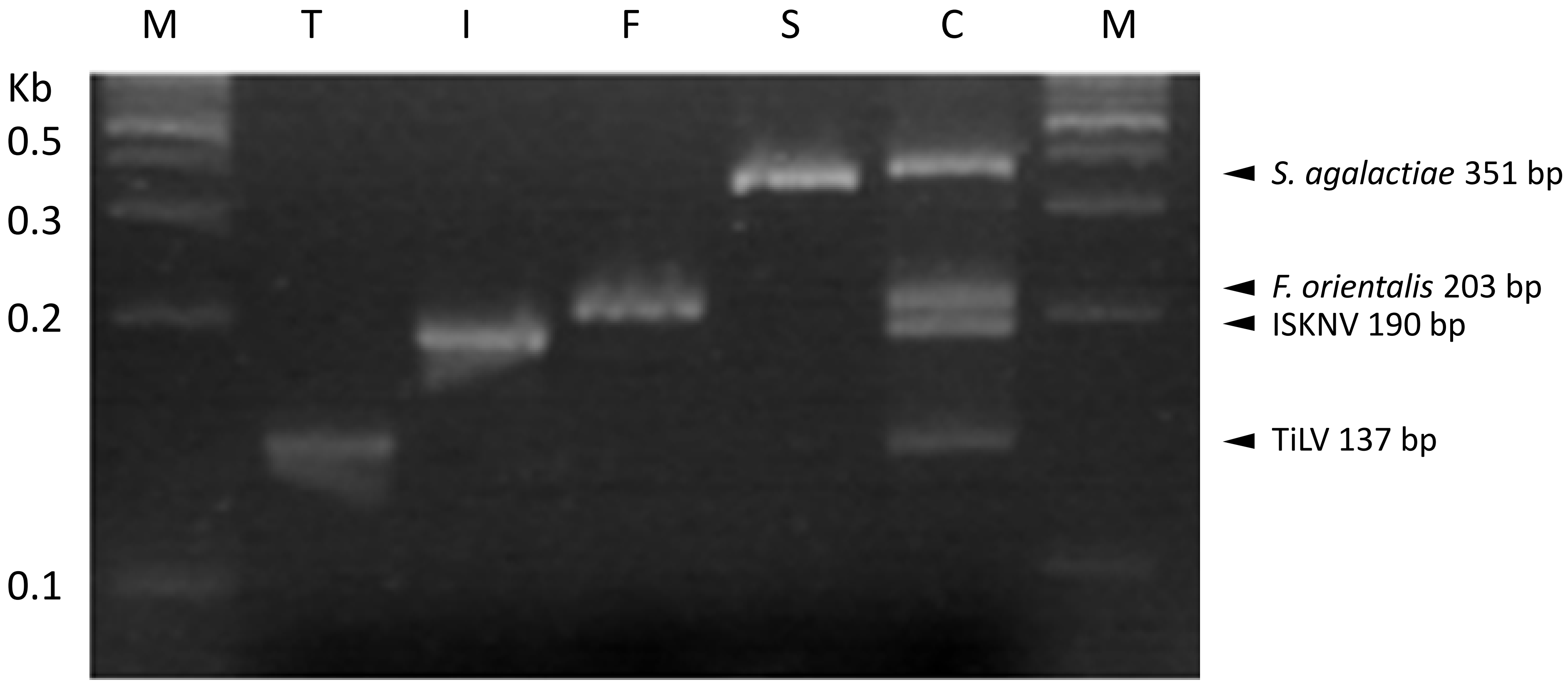

### Supplemental Figure 2

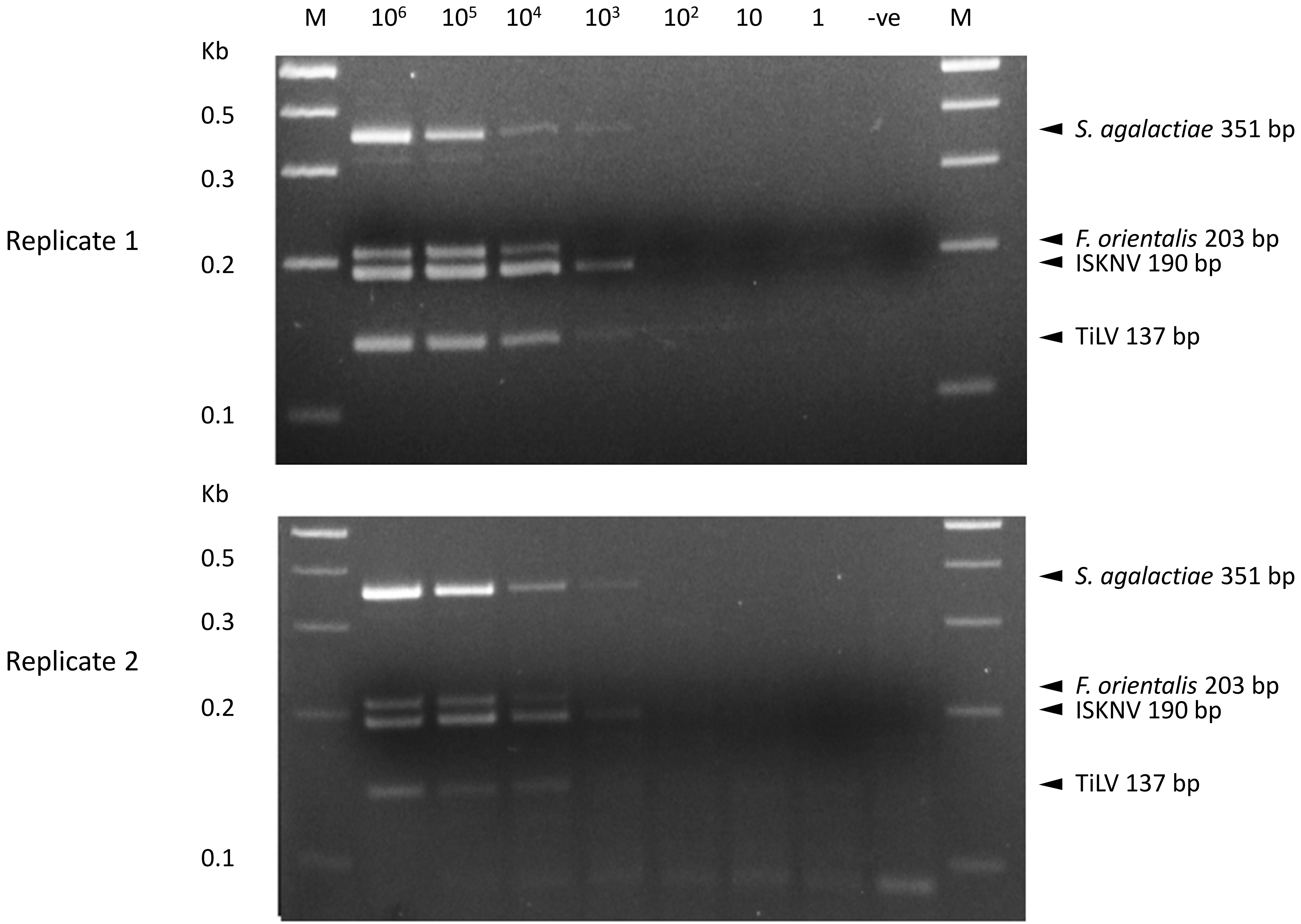
