## Supplemental Tables for "Multiplex PCR with Nanopore Sequencing for Sequence-Based Detection of Four Tilapia Pathogens"

**Supplemental Table 1.** Primers used in this experiment.

| **Target** | **Sequence (5’→3’)** | **Tm**  **(°C)** | **Size**  **(bp)** | **Reference** |
| --- | --- | --- | --- | --- |
| TiLV Segment9 | TiLV-S9-qF;  CTAGACAATGTTTTCGATCCAG  TiLV-S9-qR;  TTCTGTGTCAGTAATCTTGACAG | 54.9  56.2 | 137 | [1] |
| ISKNV - major capsid protein gene (*MCP*) | Meg-MCP160F;  TCAAAACAGACTGGCCATGC  Meg-MCP349R;  TAAATGACACCGACACCTCCTC | 59.0  59.7 | 190 | [2] |
| *F. orientalis -*hypothetical protein (*HP*) gene | Francis-n-o-F1;  GGCGTAACTCCTTTTAGCTTCC  Francis-n-o-R1;  TTAGAGGAGCTTGGAAAAGCA | 59.3  57.2 | 203 | [3] |
| *S. agalactiae - groEL* gene | SagroEL2 F;  GCAAGTTTTAGGACAGTCTGCT  SagroEL2 R;  AGTTTCAGTGCCGCTACTTT | 58.9  57.7 | 351 | [4] |

Tm, melting temperature, bp, base pair.

**Supplemental Table 2.** Sources of plasmid template used in sensitivity assay.

| Pathogen | Description | Reference |
| --- | --- | --- |
| TiLV | Segment 9 (351 bp)/pGEM-T easy | [5] |
| ISKNV | MCP (1,362 bp)/pGEM-T easy | [6] |
| *F. orientalis* | HP (203 bp)/pPerfect | [3] |
| *S. agalactiae* (SAG) | groEL (351 bp)/pGEM-T easy | This study |

**Supplemental Table 3.** Reaction mixture and cycling conditions for multiplex PCR diagnosis.

| Reagent | Volume (µL) | Final conc. |
| --- | --- | --- |
| Template | 4 | Up to 400 ng |
| 10 µM groEL F | 0.4 | 160 nM |
| 10 µM groEL R | 0.4 | 160 nM |
| 10 µM Francis-n-oF1 | 0.4 | 160 nM |
| 10 µM Francis-n-oR1 | 0.4 | 160 nM |
| 10 µM TiLV-S9-qF | 0.2 | 80 nM |
| 10 µM TiLV-S9-qR | 0.2 | 80 nM |
| 10 µM Meg-MCP160 | 0.2 | 80 nM |
| 10 µM Meg-MCP349R | 0.2 | 80 nM |
| 2X KAPA SYBR FAST qPCR Master Mix | 12.5 | 1X |
| 50X KAPA RT Mix | 0.5 | 1X |
| 10 mM dNTPs | 0.4 | 160 µM |
| D.W. | 5.2 |  |
| Total | 25 |  |

| Step | Condition | Cycle |
| --- | --- | --- |
| Reverse transcription (RT) | 42°C 5 min | 1 |
| RT inactivation | 95°C 3 min | 1 |
| Denaturation | 95°C 10 sec | 40 |
| Annealing and extension | 60°C 30 sec |  |
| Final extension | 60°C 5 min | 1 |
| Hold | 12°C $\infty$ | - |

**Supplemental Table 4.** Summary of qPCR testing for the detection and quantification of pathogens in tilapia tissue samples

| **Pathogen** | **Method** | **Target** | **Reaction** | **Cycling conditions** | **qPCR performance** | **Reference** |
| --- | --- | --- | --- | --- | --- | --- |
| **Tilapia lake virus (TiLV)** | Hydrolysis probe RT-qPCR | TiLV Segment9 | A 20 µL qPCR reaction contained the 200 ng template, 450 nM of each primer, 150 nM probe and 1X qScript One-Step RT-qPCR Kit (Quanta bio, Cat no#95134-500) | Reverse transcription 50°C for 10 min, 95°C for 1 min and 40 cycles of 95°C for 10 s and 58°C for 30 s | Y = -3.476 X + 42.295  R^2^ = 0.998  E = 94.0% | [1] |
| **Infectious spleen and kidney necrosis virus (ISKNV)** | Hydrolysis probe qPCR | ISKNV major capsid protein gene (MCP) | A 20 µL qPCR reaction contained the 200 ng template, 900 nM of each primer, 250 nM probe and 1X iTaq Universal Probes supermix (BioRAD, Cat no#172-5131) | 95°C for 10 min and 40 cycles of 95°C for 15 s and 64°C for 1 min | Y = -3.480X + 41.674  R^2^ = 0.997  E = 93.8% | [7] |
| ***Francisella orientalis* (*FnO*)** | SYBR qPCR | *F. orientalis* hypothetical protein (HP) gene | A 20 µL qPCR reaction contained the 200 ng template, 200 nM of each primer and 1X KAPA SYBR FAST qPCR Master Mix  (KapaBiosystems, Cat no#KK4600) | 95°C for 3 min and 40 cycles of 95°C for 30 s and 60°C for 30 s followed by melt curve analysis | Y = -3.563X + 39.873  R^2^ = 0.999  E = 90.8% | This study |
| ***Streptococcus agalactiae* (SAG)** | Hydrolysis probe qPCR | *S. agalactiae* groEL gene | A 20 µL qPCR reaction contained the 300 ng template, 900 nM of each primer, 250 nM probe and 1X iTaq Universal Probes supermix (BioRAD, Cat no#172-5131) | 95°C for 2 min and 40 cycles of 95°C for 5 s and 60°C for 30 s | Y = -3.581X + 43.424  R^2^ = 0.991  E = 90.2% | Modified from  [4] |

**Supplemental Table 5**. Sequencing and alignment statistics

| Sample | Raw Reads | Primer Trimmed | Aligned | Aligned Read Distribution | | | | Minimap2 of previously unaligned raw reads (default setting) | | | | | BlastN of remaining unaligned reads  (-word_size=15, evalue=0.01) | | | | |
| --- | --- | --- | --- | --- | --- | --- | --- | --- | --- | --- | --- | --- | --- | --- | --- | --- | --- |
|  |  |  |  | FnO | ISKNV | SAG | TILV | Tilapia | FnO | ISKNV | SAG | TILV | Tilapia | FnO | ISKNV | SAG | TILV |
| r1_4PAT_1 | 9030 | 3543 | 2989 | 1329 | 669 | 770 | 221 | 0 | 1763 | 769 | 1589 | 104 | 11 | 567 | 317 | 210 | 333 |
| r1_4PAT_DIL_1 | 22933 | 10659 | 9119 | 3825 | 1838 | 2970 | 486 | 0 | 3290 | 1306 | 4724 | 132 | 41 | 1298 | 744 | 678 | 613 |
| r1_FNO_1 | 13692 | 5913 | 5265 | 5239 | 8 | 16 | 2 | 0 | 6000 | 5 | 41 | 0 | 32 | 1932 | 3 | 8 | 0 |
| r1_ISKNV_1 | 10103 | 5440 | 4464 | 10 | 4443 | 10 | 1 | 0 | 8 | 2783 | 50 | 0 | 11 | 4 | 2014 | 18 | 1 |
| r1_SAG_1 | 70735 | 31116 | 29590 | 39 | 22 | 29528 | 1 | 0 | 25 | 5 | 32940 | 0 | 80 | 7 | 5 | 6043 | 2 |
| r1_TILV_1 | 32 | 9 | 3 | 0 | 0 | 0 | 3 | 0 | 2 | 3 | 5 | 0 | 0 | 1 | 2 | 4 | 6 |
| r2_NT_F1 | 4262 | 1762 | 671 | 72 | 10 | 0 | 589 | 524 | 131 | 23 | 2 | 263 | 385 | 50 | 13 | 0 | 1533 |
| r2_NT_F2 | 6827 | 2720 | 881 | 120 | 36 | 0 | 725 | 611 | 256 | 54 | 0 | 335 | 655 | 141 | 37 | 1 | 2504 |
| r2_NT_F3 | 10725 | 5347 | 1832 | 108 | 34 | 0 | 1690 | 1066 | 222 | 59 | 1 | 690 | 844 | 101 | 30 | 0 | 4320 |
| r2_NT_F4 | 16668 | 8340 | 2920 | 59 | 16 | 0 | 2845 | 757 | 151 | 24 | 0 | 1287 | 707 | 83 | 18 | 0 | 7440 |
| r2_NT_F5 | 15823 | 7414 | 2572 | 76 | 21 | 0 | 2475 | 1315 | 208 | 47 | 0 | 1188 | 1392 | 101 | 18 | 1 | 6516 |
| r2_NT_Gh23 | 4316 | 1374 | 886 | 37 | 836 | 0 | 13 | 741 | 51 | 934 | 0 | 5 | 713 | 24 | 539 | 0 | 18 |
| r2_NT_Gh24 | 4033 | 629 | 151 | 71 | 56 | 0 | 24 | 1780 | 141 | 67 | 0 | 11 | 1269 | 53 | 20 | 0 | 35 |
| r2_NT_Gh25 | 7663 | 1081 | 601 | 144 | 417 | 5 | 35 | 3560 | 251 | 392 | 6 | 14 | 1676 | 137 | 237 | 0 | 51 |
| r2_RT_EX03 | 8167 | 3045 | 2663 | 2632 | 18 | 0 | 13 | 179 | 3452 | 12 | 0 | 11 | 88 | 1227 | 9 | 0 | 36 |
| r2_RT_FM1A | 14061 | 5156 | 4491 | 4422 | 32 | 0 | 37 | 67 | 5712 | 16 | 0 | 10 | 102 | 2387 | 11 | 1 | 54 |
| r2_RT_FM1B | 7375 | 2038 | 1751 | 1728 | 9 | 3 | 11 | 36 | 3666 | 5 | 5 | 5 | 34 | 1267 | 6 | 3 | 21 |
| r2_RT_M1 | 8861 | 2968 | 2542 | 2512 | 16 | 0 | 14 | 26 | 3727 | 12 | 0 | 7 | 77 | 1667 | 9 | 0 | 16 |
| r3_NT_GroupB | 1638 | 620 | 594 | 1 | 2 | 590 | 1 | 29 | 0 | 1 | 635 | 0 | 16 | 3 | 0 | 190 | 1 |
| r3_NT_ZoneA | 2258 | 531 | 505 | 2 | 2 | 499 | 2 | 11 | 3 | 1 | 715 | 0 | 26 | 2 | 1 | 213 | 3 |
| r3_NT_ZoneB | 427 | 54 | 38 | 4 | 1 | 30 | 3 | 189 | 6 | 1 | 44 | 1 | 51 | 3 | 0 | 26 | 0 |
| r3_SAG05 | 334 | 69 | 57 | 2 | 1 | 53 | 1 | 4 | 8 | 1 | 66 | 0 | 4 | 8 | 1 | 59 | 4 |
| r3_SAG15 | 561 | 176 | 163 | 2 | 2 | 158 | 1 | 2 | 7 | 1 | 178 | 2 | 10 | 4 | 0 | 86 | 3 |
| r3_SAG30 | 901 | 391 | 370 | 2 | 2 | 365 | 1 | 1 | 8 | 0 | 333 | 0 | 5 | 0 | 0 | 98 | 1 |
| r3_SAG50 | 1662 | 624 | 584 | 6 | 3 | 573 | 2 | 3 | 15 | 2 | 535 | 1 | 8 | 3 | 0 | 234 | 5 |
| r3_TILV05 | 166 | 37 | 25 | 4 | 2 | 15 | 4 | 6 | 8 | 0 | 35 | 5 | 3 | 5 | 0 | 15 | 13 |
| r3_TILV15 | 357 | 158 | 76 | 4 | 0 | 9 | 63 | 5 | 5 | 0 | 20 | 29 | 4 | 1 | 1 | 7 | 108 |
| r3_TILV30 | 357 | 140 | 83 | 4 | 1 | 19 | 59 | 4 | 6 | 3 | 42 | 29 | 2 | 0 | 2 | 20 | 85 |
| r3_TILV50 | 1309 | 675 | 345 | 2 | 1 | 28 | 314 | 5 | 10 | 0 | 43 | 66 | 6 | 6 | 0 | 30 | 455 |
| r3_4PAT_DIL_3 | 1480 | 477 | 410 | 129 | 24 | 206 | 51 | 1 | 178 | 25 | 280 | 10 | 14 | 79 | 8 | 114 | 50 |

[1]	S. Taengphu, P. Kayansamruaj, Y. Kawato, J. Delamare-Deboutteville, C.V. Mohan, H.T. Dong, S. Senapin, Concentration and quantification of Tilapia tilapinevirus from water using a simple iron flocculation coupled with probe-based RT-qPCR, PeerJ. 10 (2022) e13157. https://doi.org/10.7717/peerj.13157.

[2]	Y. Kawato, D.M. Cummins, S. Valdeter, P.G. Mohr, T. Ito, K. Mizuno, H. Kawakami, L.M. Williams, M.S.J. Crane, N.J.G. Moody, Development of New Real-time PCR Assays for Detecting *Megalocytivirus* Across Multiple Genotypes, Fish Pathology. 56 (2021) 177–186. https://doi.org/10.3147/jsfp.56.177.

[3]	H.T. Dong, W. Gangnonngiw, K. Phiwsaiya, W. Charoensapsri, V.V. Nguyen, P. Nilsen, P.J. Pradeep, B. Withyachumnarnkul, S. Senapin, C. Rodkhum, Duplex PCR assay and in situ hybridization for detection of Francisella spp. and Francisella noatunensis subsp. orientalis in red tilapia, Dis Aquat Organ. 120 (2016) 39–47. https://doi.org/10.3354/dao03021.

[4]	W.J. Leigh, R.N. Zadoks, A. Jaglarz, J.Z. Costa, G. Foster, K.D. Thompson, Evaluation of PCR primers targeting the groEL gene for the specific detection of Streptococcus agalactiae in the context of aquaculture, J Appl Microbiol. 125 (2018) 666–674. https://doi.org/10.1111/jam.13925.

[5]	Y. Thawornwattana, H.T. Dong, K. Phiwsaiya, P. Sangsuriya, S. Senapin, P. Aiewsakun, Tilapia lake virus (TiLV): Genomic epidemiology and its early origin, Transboundary and Emerging Diseases. 68 (2021) 435–444. https://doi.org/10.1111/tbed.13693.

[6]	H.T. Dong, S. Jitrakorn, P. Kayansamruaj, N. Pirarat, C. Rodkhum, T. Rattanarojpong, S. Senapin, V. Saksmerprome, Infectious spleen and kidney necrosis disease (ISKND) outbreaks in farmed barramundi (Lates calcarifer) in Vietnam, Fish Shellfish Immunol. 68 (2017) 65–73. https://doi.org/10.1016/j.fsi.2017.06.054.

[7]	Y. Kawato, P.G. Mohr, M.S.J. Crane, L.M. Williams, M.J. Neave, D.M. Cummins, M. Dearnley, S. Crameri, C. Holmes, J. Hoad, N.J.G. Moody, Isolation and characterisation of an ISKNV-genotype megalocytivirus from imported angelfish Pterophyllum scalare, Diseases of Aquatic Organisms. 140 (2020) 129–141. https://doi.org/10.3354/dao03499.
